# Store-Operated Ca^2+^ Channels Mediate Microdomain Ca^2+^ Signals and Amplify Gq-Coupled Ca^2+^ Elevations in Capillary Pericytes

**DOI:** 10.1101/2022.05.25.493438

**Authors:** Braxton Phillips, Jenna Clark, Éric Martineau, Ravi L. Rungta

**Affiliations:** Department of Neuroscience, Université de Montréal, Montréal, Québec, Canada; Department of Physiology and Pharmacology, Université de Montréal, Montréal, Québec, Canada; Department of Stomatology, Faculty of Dental Medicine, Université de Montréal, Montréal, Québec H3C3J7, Canada; Centre interdisciplinaire de recherche sur le cerveau et l’apprentissage, Université de Montréal, Montréal, Québec, Canada

## Abstract

Pericytes are multifunctional cells of the vasculature that are vital to brain homeostasis, yet many of their fundamental physiological properties remain unexplored. Ca^2+^ is a ubiquitous second messenger across cell-types, where it mediates diverse functions such as contractility and gene transcription. It is therefore important to understand the cellular mechanisms underlying Ca^2+^ signalling in capillary pericytes. Here, we performed pharmacological and ion substitution experiments to investigate the mechanisms underlying pericyte Ca^2+^ signaling in acute cortical brain slices of PDGFRβ-GCaMP6f mice. We report that in mid-capillary bed pericytes (≥ 4^th^ branch order), spontaneous microdomain Ca^2+^ signals are dependent on extracellular Ca^2+^, but largely independent of depolarization, L- and T-type voltage-gated calcium channels (VGCCs), and TRPC3/6 channels. In contrast, these microdomain signals were inhibited by multiple Orai channel blockers, including the specific antagonist GSK-7975A. Furthermore, capillary pericytes exhibited classical store operated calcium entry (SOCE) following store depletion that was sensitive to GSK-7975A and required for amplification of intracellular Ca^2+^ increases evoked by the vasoconstrictor endothelin-1. These results suggest that Orai SOCE mediates microdomain Ca^2+^ signals at rest and amplifies Gq-GPCR coupled Ca^2+^ elevations in capillary pericytes. Thus, SOCE is a major regulator of pericyte Ca^2+^ and a target for manipulating their function in health and disease.

## Introduction

The mural cells of the brain are multifunctional cells organized in a continuum along the arterio-venous axis of the cerebral vasculature. This continuum is defined by distinct cell types, which includes smooth muscle cells (SMCs) on penetrating arterioles, pre-capillary sphincters and ensheathing pericytes (also named terminal SMCs) of the arteriole-to-capillary transition zone, capillary pericytes of the mid-capillary bed (of mesh and thin-strand morphology), and venule SMCs (Grubb et al., 2021; Hartmann et al., 2022). Although they share a common origin (Armulik et al., 2011) and similar nomenclature, these cell types have numerous morphological (Grant et al., 2017; Ratelade et al., 2020), transcriptomic (Vanlandewijck et al., 2018; Yang et al., 2022), and functional differences (Hartmann et al., 2021; Rungta et al., 2018). For example, SMCs and ensheathing pericytes fully encircle and exhibit near complete coverage of the endothelial tube, highly express alpha smooth muscle actin (α-SMA), and undoubtedly control the regulation of cerebral blood flow (CBF). Capillary pericytes have thin or mesh-like processes that run longitudinal to the vessel, express little to no α-SMA, and their role in controlling CBF in physiological contexts remains poorly defined. Nevertheless, blood flow control aside, capillary pericytes have numerous functions in health and disease, such as blood-brain-barrier regulation (Armulik et al., 2010; Bell et al., 2010; Daneman et al., 2010) neuroimmune regulation (Rustenhoven et al., 2017), angiogenesis (Gerhardt and Betsholtz, 2003), glial scar formation (Dias et al., 2021), and suggested stem cell-like properties (Nakagomi et al., 2015), which may all be affected by second messengers such as Ca^2+^. Yet, despite these known functions, the basic physiological properties of brain capillary pericytes, including the mechanisms that regulate their Ca^2+^ signaling, remain understudied.

All mural cells exhibit spontaneous fluctuations in intracellular Ca^2+^, which are termed Ca^2+^ transients (Glück et al., 2021; Hill et al., 2015; Rungta et al., 2018). In SMCs and ensheathing pericytes, Ca^2+^ transients are caused by transmembrane influx through voltage-gated Ca^2+^ channels (VGCCs) that recruits the Ca^2+^ required for α-SMA-mediated contraction (Gonzales et al., 2020; Hill-Eubanks et al., 2011; Korte et al., 2022). Capillary pericytes also exhibit Ca^2+^ transients with similar properties *in vivo* and *in vitro*, which are confined to spatial microdomains and modulated by neuronal activity (Alarcon-Martinez et al., 2020; Glück et al., 2021; Rungta et al., 2018), leading to the belief that their Ca^2+^ transients are likewise voltage dependent. Interestingly, a recent study reported that in resting conditions capillary pericyte Ca^2+^ transients were only minimally sensitive to the potent L-type VGCC blocker nimodipine but were largely blocked with the non-selective ion channel blocker SKF-96365 (Glück et al., 2021), leading to speculation of Ca^2+^ entry via pericyte TRPC channels. However, SKF-96365 blocks several other Ca^2+^ channels that are highly expressed in capillary pericytes such as T-type VGCCs (Singh et al., 2010), and the Orai family of store-operated Ca^2+^ channels (SOCE) (Várnai et al., 2009). Capillary pericytes also express a plethora of Gq-coupled receptors (Hariharan et al., 2020; Vanlandewijck et al., 2018), which when activated would be expected to trigger Ca^2+^ release from internal stores via IP_3_ receptor signaling. Therefore, a more detailed investigation is required to understand the cellular mechanisms underlying microdomain Ca^2+^ signaling in capillary pericytes, their voltage dependence, and the interplay between Ca^2+^ release from internal stores and transmembrane influx.

Here, we utilized pharmacology experiments on acute cortical brain slices from transgenic mice expressing a fluorescent calcium indicator, GCaMP6f, in brain mural cells under the PDGFRβ promoter (PDGFRβ-Cre;GCaMP6f mice) to delineate the mechanism underlying Ca^2+^ signaling in brain capillary pericytes. Surprisingly, we find that capillary pericyte Ca^2+^ signals are largely unaffected by manipulations known to alter pericyte membrane potential, or by direct inhibition of VGCCs. We show that capillary pericyte Ca^2+^ transients are, instead, primarily mediated by store operated Ca^2+^ entry (SOCE) channels fitting the pharmacological profile of Orai1/3. Underscoring the potential importance of pericyte SOCE channels, we demonstrate that they amplify Gq-coupled [Ca^2+^]_i_ elevations evoked by the potent endogenous vasoconstrictor endothelin-1 (ET-1).

## Results

Pericyte Ca^2+^ signals were visualized with confocal imaging in acute cortical slices from PDGFRβ-Cre;GCaMP6f mice. The capillary lumen was labeled with an intravenous (I.V.) injection of Rhodamine-Dextran (70kDa) and pericyte somas were labelled with the recently described pericyte specific dye TO-PRO-3 (Mai-Morente et al., 2021) (**Fig. 1A**). All capillary pericytes were ≥ 4 vascular branch segments downstream of the penetrating arteriole. The spontaneous and spatially incoherent nature of pericyte Ca^2+^ transients lead us to take advantage of the recently developed event-based analysis tool AQuA, which was developed precisely for unbiased analysis of such signals, but in astrocytes (Wang et al., 2019). When applied to pericytes, AQuA was found suitable for extracting several properties of these pericyte Ca^2+^ transients in an unbiased way, such as overall event frequency, amplitude, area, and duration (**Fig. S1; Movie S1; Table S1-S4**). Indeed, Ca^2+^ activity levels measured in manually drawn ROIs were well matched to the density of events measured in these same processes using AQuA (**Fig. 1B-D**).

**Figure 1:**
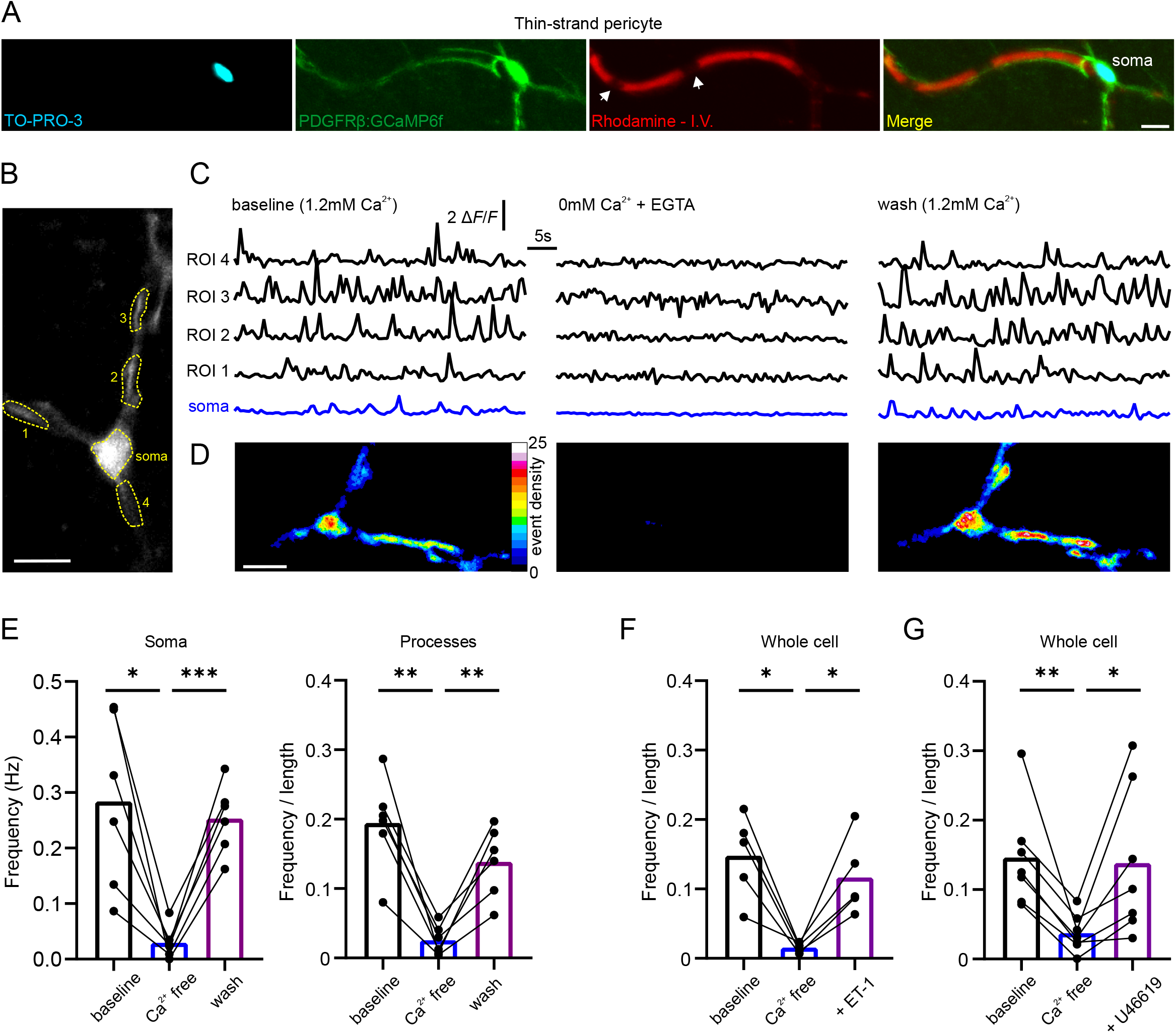
Imaging of pericyte microdomain Ca^2+^ signals and their dependence on extracellular Ca^2+^. **(A)** Confocal z-projection of a pericyte expressing GCaMP6f (green) in an acute brain slice loaded TO-PRO-3 (cyan) and contacting a capillary whose lumen is labeled with Rhodamine-dextran (Red) – arrow heads indicate red blood cells. **(B)** Image of GCaMP6f fluorescent pericyte and ROIs selected for **(C)**. **(C)** ROI based analysis of fluorescent changes in ROIs from **(B)** at baseline, in Ca^2+^ free ACSF, and following wash back of Ca^2+^ containing ACSF. **(D)** Heat maps showing event density calculated from AQuA (percent time each pixel is active), at baseline, in Ca^2+^ free ACSF, and following wash back of Ca^2+^ containing ACSF. **(E)** Summarized data showing effect of transiently removing and washing back Ca^2+^ on pericyte Ca^2+^ transient frequency in the soma and processes. Statistics were calculated by repeated measures one-way ANOVA followed by Tukey’s multiple comparisons test, baseline and wash were not significantly different. **(F, G)** 10 nm ET-1 **(F)**, or 100 nm U46619 **(G)** increases Ca^2+^ transient frequency in 0 mM extracellular Ca^2+^ ACSF. Statistics were calculated by repeated measures one-way ANOVA followed by Šídák’s multiple comparisons test. All n values and p-values can be found in Supplemental Tables 1 and 2. All image scale bars, 10 µm.

We first tested whether these microdomain pericyte microdomain calcium transients depended on extracellular Ca^2+^ by washing out extracellular Ca^2+^ from the perfused artificial cerebral spinal fluid (ACSF). Indeed, perfusion of 0 mM Ca^2+^, 2 mM EGTA solution rapidly and reversibly depressed the frequency, size, and duration of Ca^2+^ transients in both the processes and soma of the pericyte (**Fig. 1D-E; Table S1**). To confirm that intracellular store Ca^2+^ was not completely depleted in Ca^2+^ free ACSF, we applied Gq-coupled GPCR agonists for either the endothelin (ET)-A or thromboxane A2 receptor, ET-1 and U46619, which increases intracellular pericyte-Ca^2+^ in an IP_3_-dependent manner (Glück et al., 2021). ET-1 and U46619 were still able to transiently evoke Ca^2+^ signals (**Fig. 1E,F, Table S2**), suggesting that spontaneous calcium transients in capillary pericytes depend on transmembrane influx rather than internal stores. Intriguingly, ET-1 and U46619 induced elevations in Ca^2+^ transient frequency rather than a sustained and global elevation, and this increase in transient frequency was temporary (ET-1, 248.2 +/- 46.52 seconds; U46619, 268.6 +/- 32.65 seconds), hinting at the necessity of extracellular Ca^2+^ influx to refill internal stores following their depletion by IP_3_R mediated release.

In SMCs and ensheathing pericytes, depolarization triggers transmembrane influx of Ca^2+^ via VGCCs, thereby increasing Ca^2+^ transient frequency and [Ca^2+^]_i_ (Gonzales et al., 2020; Hill-Eubanks et al., 2011; Korte et al., 2022). Therefore, we tested the effect of blocking the predominantly expressed pericyte VGCC subtypes, Ca_V_1.2 (L-type) and Ca_V_3.2 (T-type), with nifedipine and Z944, respectively (**Fig. 2A**). Application of nifedipine (20 µM) and Z944 (2 µM) had no effect on the frequency of calcium transients, or baseline calcium levels, in either the processes or soma (**Fig. 2B, Table S3,S4**). However, we could not exclude the possibility that VGCC transients would become more apparent if the membrane were depolarized, as previously reported for ensheathing type pericytes (Gonzales et al., 2020). Surprisingly, depolarization by increasing extracellular K^+^ concentration from 2.5 mM to 60 mM did not increase thin-strand/mesh pericyte Ca^2+^ transient frequency or baseline Ca^2+^ levels in either the processes or soma (**Fig. 2C, Table S3,S4**). Furthermore, VGCC blockers still had no effect on thin-strand pericyte Ca^2+^ in 60 mM K^+^ solution (**Fig. 2D-F, Table S3,S4**). This contrasts with the effect of VGCCs on ensheathing-type pericyte Ca^2+^, whose transient frequency in 60 mM K^+^ was significantly reduced in the presence of VGCC blockers (**Fig. 2D-F, Table S3,S4**, 1^st^ to 3^rd^ branch order from penetrating arteriole). It is worth noting that a brief decrease in capillary pericyte Ca^2+^ transient frequency was observed upon application of K^+^, which recovered slightly and stabilized at levels below but not significantly different from baseline (**Fig. 2C**). Finally, we set out to test the effect of pharmacologically altering K_ATP_ channel activity, whose modulation has been shown to have large effects on pericyte membrane potential (Hariharan et al., 2022; Li and Puro, 2001; Sancho et al., 2022). Consistent with our above results showing a lack of depolarization induced Ca^2+^ influx in capillary pericytes, application of pinacidil (10 µM), a K_ATP_ channel opener which hyperpolarizes pericytes, had no effect on transient frequency in processes or the soma (**Fig. 2G**). Similarly, glibenclamide (20 µM) a non-selective K_ATP_ channel blocker, decreased rather than increased transient frequency in pericyte processes (**Fig. 2H**). These results suggest that, in contrast to SMCs and ensheathing pericytes which exhibit increased Ca^2+^ entry upon depolarization via VGCCs, capillary pericyte Ca^2+^ transients are mostly independent of VGCCs and membrane potential.

**Figure 2:**
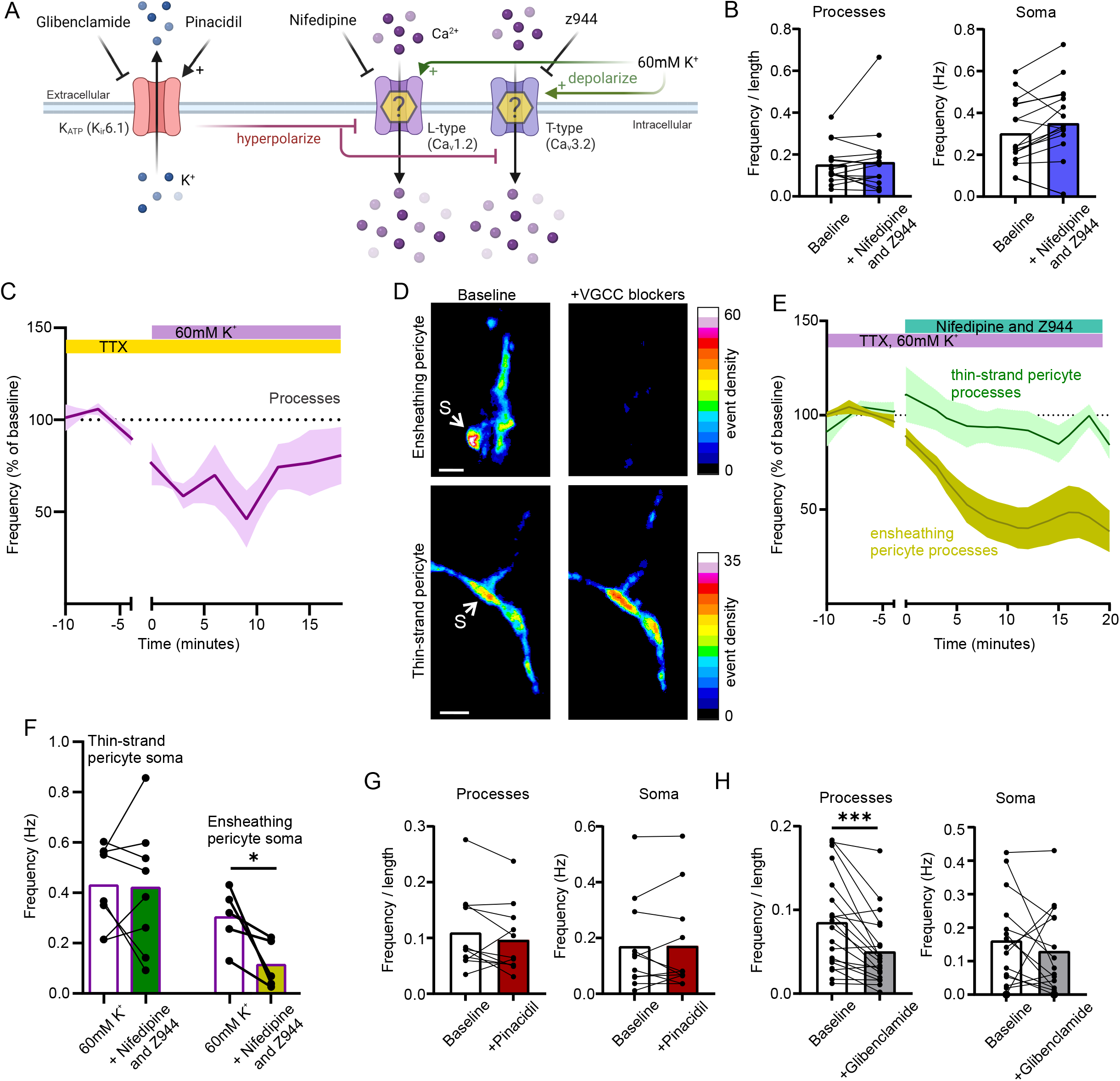
Pericyte Ca^2+^ transients are not potentiated by depolarization or mediated by VGCCs. **(A)** Schematic of experimental plan in Fig. 2. Created with biorender.com **(B)** Blocking L-type and T-type VGCCs with 20 µM nifedipine and 2 µM Z944 did not reduce the frequency of Ca^2+^ transients in capillary pericyte processes or soma. **(C)** Changing extracellular [K^+^] from 2.5 to 60 mM does not increase the frequency of Ca^2+^ transients in capillary pericytes (data shown is from processes). **(D)** Example images showing event density (percentage of frames an event was detected within each pixel) after application of 20 µM nifedipine and 2 µM Z944 in 60 mM K^+^ ACSF, for an ensheathing pericyte (3^rd^ order, top), and a thin-strand pericyte (bottom). S points to soma. **(E)** Summarized data, showing application of 20 µM nifedipine and 2 µM Z944 in 60 mM K^+^ ACSF reduces Ca^2+^ transient frequency in ensheathing, but not thin-strand pericyte processes and **(F)** soma. **G)** Summarized data of K_ATP_ channel opener 10 µM pinacidil on capillary pericyte Ca^2+^ transient frequency in processes, and soma. **(H)** Summarized data of K_ATP_ channel blocker 20 µM glibenclamide on capillary pericyte Ca^2+^ transient frequency in processes, and soma. All experiments performed in 500 nM TTX. Shaded area represents SEM. Time 0 represents the start of the first acquisition in the presence of the new bathing solution (when the solution is estimated to have reached the slice chamber). All n values and p-values can be found in Table S1. All image scale bars, 10 µm.

Having established a striking difference in the mechanisms of Ca^2+^ signalling between ensheathing and capillary pericytes, we next examined the unidentified calcium influx pathway(s) underlying microdomain calcium transients in capillary pericytes (**Fig. 3A**). We first tested the non-specific ion channel blocker SKF-96365, which was previously reported to block capillary pericyte Ca^2+^ transients (Glück et al., 2021). Consistent with this previous report, Ca^2+^ transient frequency in capillary pericyte processes and soma was largely diminished in the presence of SKF-96365 (100 µM) (**Fig. 3B, Table S3,S4**). As SKF-96365 is a potent TRPC channel blocker, and TRPC3/6 are highly expressed in pericytes (Glück et al., 2021; Vanlandewijck et al., 2018) and exhibit a mixed cation conductance, we further examined the sensitivity of pericyte Ca^2+^ transients to the TRPC3 blocker, Pyr3 and the potent TRPC3/6 blocker GSK-2833503A. Capillary pericyte Ca^2+^ transients were unaffected by TRPC3 inhibition with Pyr3 (20 µM), and inhibition of TRPC3/6 with GSK 2833503A (10 µM) (**Fig. 3B, Table S3,S4**), suggesting influx via another SKF-96365 sensitive pathway. Interestingly, capillary pericytes highly express Orai1 and Orai3 SOCE channels (Vanlandewijck et al., 2018), and these channels are also sensitive to SKF-96365 (Várnai et al., 2009). We therefore tested the sensitivity of capillary pericyte Ca^2+^ transients to the non-selective Orai channel blocker 2-APB. 2-APB is a potent Orai1 channel blocker at high concentration but activates Orai1 channels at low concentrations (Prakriya and Lewis, 2001). Consistent with the bidirectional concentration dependent sensitivity of Orai1 channels to 2-APB, 10 µM 2-APB increased capillary pericyte Ca^2+^ transient frequency, whereas 100 µM 2-APB nearly abolished all transients (**Fig. 3C, Table S3,S4**). Although the effects of 2-APB were strongly supportive of Orai mediated Ca^2+^ entry, 2-APB has several off-target effects, such as blocking IP_3_Rs and some TRP channels. Therefore, we tested the sensitivity of capillary pericyte Ca^2+^ transients to the Orai-specific blocker GSK-7975A (Derler et al., 2013). Indeed, GSK-7975A (40 µM) robustly reduced the frequency of Ca^2+^ transients in capillary pericytes in both the soma and processes (**Fig. 3D-G, Table S3,S4)**.

**Figure 3:**
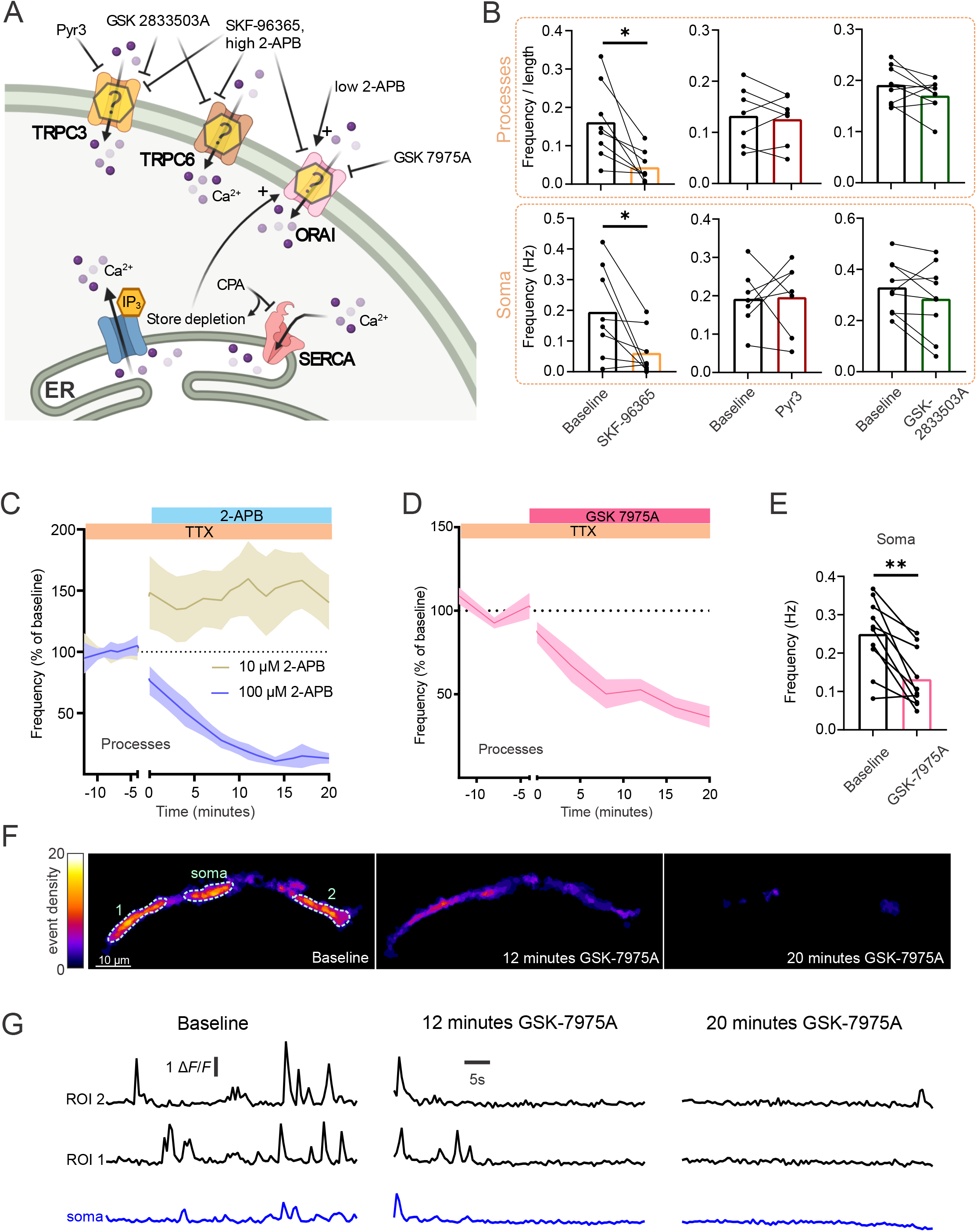
Pericyte microdomain Ca^2+^ signals are mediated by ORAI ion channels. **(A)** Schematic of experimental plan and pharmacology. Created with biorender.com **(B)** 100 µM SKF-96365 reduced the frequency of Ca^2+^ transients in capillary pericyte processes and soma. Neither the TRPC3 or the TRPC3/6 antagonists, 20 µM Pyr3 or 10 µM GSK-2833503A, significantly reduced pericyte Ca^2+^ transient frequency, soma or processes. **(C)** Low concentration (10 µM) 2-APB increases whereas high concentration (100 µM) 2-APB reduces Ca^2+^ transient frequency in capillary pericyte processes. **(D)** The Orai channel antagonist 40 µM GSK-7975A reduces Ca^2+^ transient frequency in both processes and **(E)** soma. **(F)** Event density image of a pericyte showing a decrease in Ca^2+^ transients at different time points following application of GSK-7975A **(G)** Example traces of average GCaMP6f signal over time in ROIs from soma and 2 regions of a pericytes processes in (**F**). All experiments performed in 500 nM TTX. Shaded area represents SEM. Time 0 represents the start of the first acquisition in the presence of the new bathing solution (when the solution is estimated to have reached the slice chamber). All n values and p-values can be found in Tables S3,S4.

SOCE channels link Ca^2+^ release from the endoplasmic reticulum to Ca^2+^ influx across the plasma membrane. As our data suggests that Orai channels mediate microdomain Ca^2+^ transients in capillary pericytes, we next questioned whether bona fide pericyte SOCE was also sensitive to Orai channel inhibition. A classic experiment to evoke SOCE is to deplete both endoplasmic reticulum calcium and extracellular Ca^2+^, and then to reintroduce extracellular Ca^2+^ to quantify Ca^2+^ entry via the activated SOCE channels. To perform this experiment, we applied the sarco(endo) plasmic reticulum calcium-ATPases (SERCA) inhibitor cyclopiazonic acid (CPA, 30 µM for 20 minutes) in Ca^2+^-free ACSF and then reintroduced Ca^2+^ to the bathing solution while imaging a pericyte. Re-introduction of extracellular Ca^2+^ (1.2 mM), evoked a large and prolonged increase in capillary pericyte Ca^2+^, indicative of SOCE, which was sensitive to the Orai channel blockers 2-APB and GSK-7975A (**Fig. 4A-C**). Given the above evidence supporting Orai channel mediated influx in mediating SOCE and microdomain Ca^2+^ signals in pericytes, this raised the intriguing possibility of whether Orai channels could amplify GPCR mediated Ca^2+^ elevations, following release of Ca^2+^ from internal stores, as has been reported in other cell types (e.g., (Jairaman et al., 2022)). Therefore, we tested the impact of the Orai channel blocker GSK-7975A on Ca^2+^ elevations evoked by the vasoconstrictor ET-1. Indeed, ET-1 triggered a robust and sustained Ca^2+^ elevation in pericytes (**Fig. 4D,E**), which contrasted to the temporary increase in transient frequency observed in the absence of extracellular Ca^2+^ (**Fig. 1E**). When Orai channels were blocked with GSK-7975A the magnitude of this elevation in [Ca^2+^]_i_ was strongly reduced (**Fig. 4D,E**), and as in Ca^2+^ free ACSF, ET-1 evoked a transient increase in event frequency rather than a sustained elevation (**Fig 4F,G**). These results suggest that Orai mediated SOCE amplifies Gq-GPCR mediated Ca^2+^ elevations and contributes to sustained increases in capillary pericyte Ca^2+^ following the release of vasoconstrictive agents.

**Figure 4:**
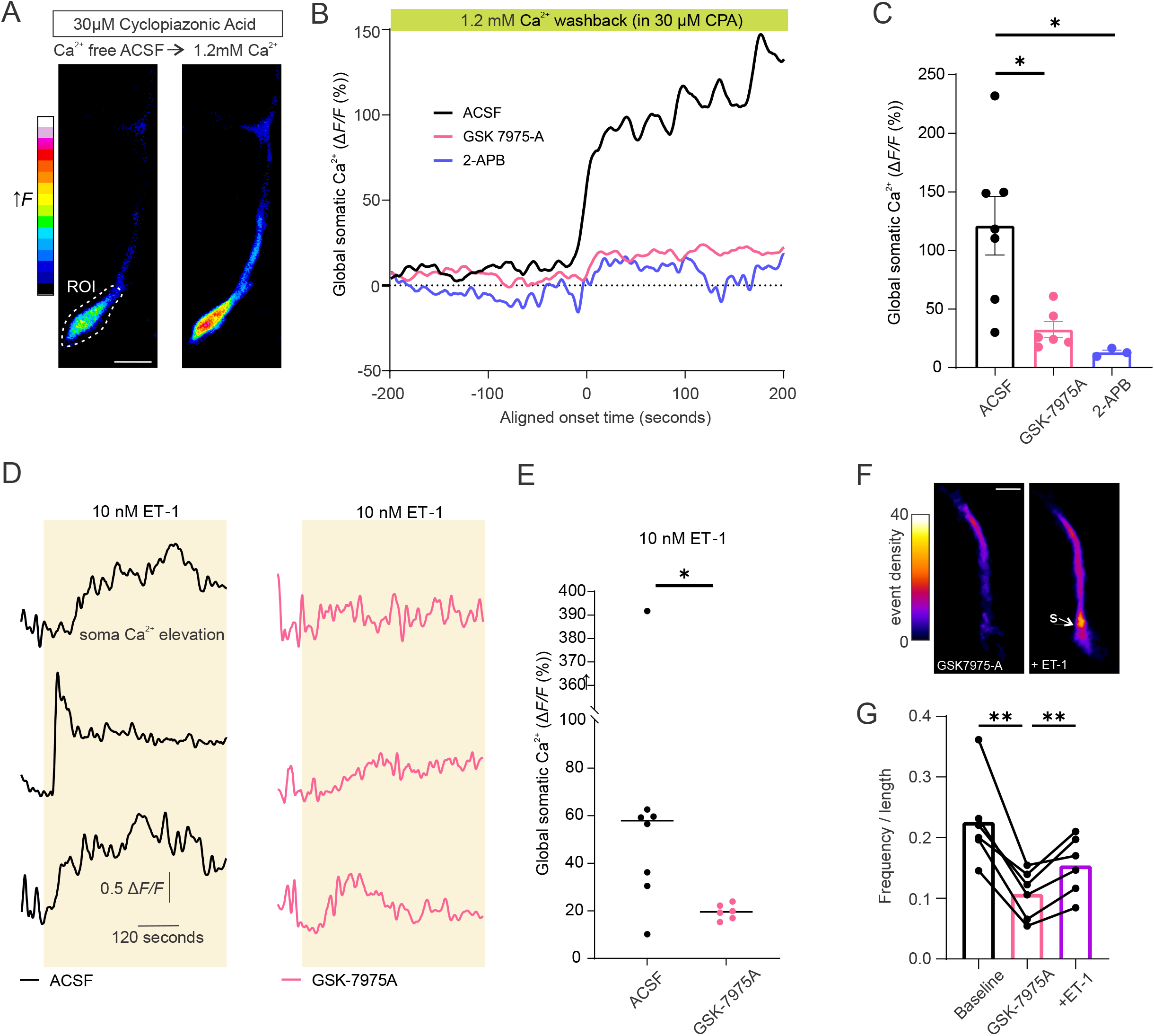
Capillary pericytes express functional SOCE that amplifies ET-1 induced [Ca^2+^] increases. **(A)** Example image showing increase in pericyte GCaMP6f fluorescence when extracellular ACSF is switched from Ca^2+^-free to 1.2mM Ca^2+^ in conditions where endoplasmic reticulum stores are depleted with 30 µM cyclopiazonic acid (CPA). An ROI around soma is used for quantification in (B) and (C). **(B)** Representative traces from example cells in which 1.2 mM Ca^2+^ is washed back on the slice in 30 µM CPA, to quantify the magnitude of Ca^2+^ entry via store operated channels, and its inhibition by 40 µM GSK-7975A and 100 µM 2-APB. **(C)** Summarized data showing effect of 40 µM GSK-7975A and 100 µM 2-APB on SOCE, following transition from 0 mM to 1.2 mM Ca^2+^ ACSF in the presence of 30 µM CPA, as in (B) and (C). Statistics were calculated with one-way ANOVA followed by Tukey’s multiple comparisons test. Error bars represent SEM. **(D)** Example traces from, showing GSK-7975A reduces the amplitude and duration of the ET-1 evoked Ca^2+^ increase. 6 cells from 6 slices (3 per condition). **(E)** Summarized data showing 40 µM GSK-7975A reduces the ET-1 evoked rise in intracellular Ca^2+^. Statistics were calculated by Mann-Whitney *U* test. Bars represent median values. **(F)** Heat map showing event density before and after ET-1 is added in the presence of GSK-7975A, analyzed with AQUA. S indicates pericyte soma. **(G)** Summarized data showing microdomain event frequency increasing temporarily, when ET-1 is added in the presence of GSK-7975A. Statistics were calculated by repeated measures one-way ANOVA followed by Šídák’s multiple comparisons test. All experiments performed in 500 nM TTX.

## Discussion

Here we demonstrate that SOCE is a major source of Ca^2+^ influx in capillary pericytes, contributing both to the generation of microdomain Ca^2+^ transients and amplification of Gq-coupled Ca^2+^ signals. Pharmacological blockade with the selective Orai channel blocker GSK-7975A and the non-selective blockers 2-APB and SKF-96365 reveal the highly Ca^2+^ selective family of Orai channels as the molecular entity underlying this Ca^2+^ entry. Consistent with these results, transcriptomic data show that pericytes express both Orai1 and 3 mRNA in rodents (Vanlandewijck et al., 2018) and in humans (Yang et al., 2022). Although we do not attempt to distinguish between ORAI isoforms in this study, the fact that Orai channels largely form hetereomers in endogenous systems (Yoast et al., 2020) suggests the molecular identity of pericyte SOCE to be Orai1/3. While TRPC channels have also been shown to interact with SOCE in certain conditions (Molnár et al., 2016; Ong et al., 2016), pericyte Ca^2+^ transients were not significantly inhibited by pharmacological inhibition of TRPC3/6 channels. However, we cannot completely rule out a potential role for TRPC1 channels in contributing to these signals.

Our work is largely in agreement with a previous study which found that Ca^2+^ transients in capillary pericytes were independent of L-type VGCCs at rest, with only soma signals being minimally sensitive to nimodipine (Glück et al., 2021). Here we extend on these results and further show that these transients are not increased by depolarization and are likewise also independent of low-voltage-activated T-type calcium channels (Ca_V_3.2), which pericytes robustly express (Vanlandewijck et al., 2018). These findings contrast with our initial hypothesis that these transients would require VGCC activity, which was based on our previous finding that [Ca^2+^]_i_ and Ca^2+^ transient frequency decreases in thin-strand pericytes following local increases in neuronal activity, although on a slower time scale than upstream ensheathing pericytes and SMCs (Rungta et al., 2018). Importantly, although our data suggests Ca^2+^ transients in capillary pericytes are largely independent of changes in membrane potential, this does not exclude a role for pericyte hyperpolarization in functional hyperemia. Indeed, recent work has demonstrated that activation of capillary pericyte K_ATP_ channels hyperpolarizes pericytes, which can then propagate this membrane hyperpolarization to upstream mural and endothelial cells (Hariharan et al., 2022; Sancho et al., 2022). Retrograde hyperpolarization is a robust phenomenon in the microvasculature and rapidly closes VGCCs on upstream SMCs and ensheathing pericytes, thereby decreasing intracellular Ca^2+^ and dilating these contractile cells to increase local cerebral blood flow (Chen et al., 2014; Iadecola et al., 1997; Longden et al., 2017; Rungta et al., 2018). Interestingly we found that blocking K_ATP_ channels with a non-specific antagonist, glibenclamide led to a decrease in the frequency of Ca^2+^ transients, as did depolarization with 60 mM K^+^ solution, consistent with a decreased driving force for Orai mediated Ca^2+^ influx. However, it cannot be ruled out that these manipulations depolarize other neighboring cells, such as astrocytes or endothelial cells, which then release factors to depress Ca^2+^ signals in the pericytes.

It is now well appreciated that the mural cells of the cerebral vasculature represent a continuum, with morphological, molecular, and functional diversity along the arterio-venous axis. During neurovascular coupling ensheathing pericytes of the capillary-arteriole transition zone (which robustly express α-SMA) dilate rapidly, whereas the capillaries of the mid-capillary bed, which are contacted by thin strand pericytes (low or negative for α-SMA) increase their diameter more slowly (Rungta et al., 2021, 2018). It remains to be determined whether this capillary dilation is a purely passive process or has an active component mediated by the pericyte. Interestingly, under pathological conditions such as Alzheimer’s disease and stroke, pericytes of the mid-capillary bed are found to be constricted (Korte et al., 2022; Nortley et al., 2019), and prolonged optogenetic activation of ChR2 on capillary pericytes has been shown to locally constrict the capillaries that they contact on a slow timescale (Hartmann et al., 2021). Here we show that SOCE is required to amplify Ca^2+^ entry mediated by ET-1. ET-1 is an endogenous vasoconstrictor that is central to pericyte pathology, which is released in ischemia (Lampl et al., 1997), and which is elevated downstream of Amyloid-β oligomers to constrict pericytes in Alzheimer’s disease (Nortley et al., 2019). In ensheathing pericytes (1-3^rd^ branch order) ET-1 evoked [Ca^2+^]_i_ increase was also recently shown to require an amplification step, but via Ca^2+^ activated Cl^-^ channel mediated depolarization and L-type VGCC activation (Korte et al., 2022). Our results indicate a similar amplification process in capillary pericytes, but with a clear molecular divergence in the mechanisms mediating ET-1 evoked Ca^2+^ influx. As Ca^2+^ is a ubiquitous signal transduction molecule throughout biology, mediating an array of cellular functions, including gene transcription and contraction, SOCE channels may therefore play an important role in numerous pericyte functions and contribute to their dysfunction in disease.

## METHODS

### Ethics statement and animals

All procedures conformed to the guidelines of the Canadian Council on Animal Care and were approved by the “Comité de déontologie sur l’expérimentation animale” (CDEA) of the Université de Montréal (QC, Canada). PDGFRβ-Cre mice (Cuttler et al., 2011) were crossed with Ai95(RCL-GCaMP6f)-D reporter mice (Jackson Laboratory) to obtain PDGFRβ-Cre; GCaMP6f-floxed double transgenic mice. Male and female mice aged P28-P134 were used in experiments.

### Acute brain slice preparation

Prior to slicing, mice were put into deep anesthesia with isoflurane and 50 uL of Rhodamine B isothiocyanate-Dextran (70 kDa, 2.5% wt:vol, Sigma-Aldrich) was injected retro-orbitally to label the vessel lumen. Mice were euthanized by decapitation. Following extraction from the skull, brains were placed into ice-cold NMDG-based slicing solution containing (in mM): 120 N-Methyl-D-glucamine, 3 KCl, 25 NaHCO_3_, 7 MgCl_2_-6H_2_O, 1 NaH_2_PO_4_-H_2_O, 20 Glucose, 2.4 Na-pyruvate, Na-ascorbate, 1 CaCl_2_-H_2_O. Then, 300µm thick coronal cortical slices were cut with a Leica VT 1200S Vibratome. Slices were transferred to a custom chamber with artificial cerebral spinal fluid (ACSF) containing (in mM): 126 NaCl, 2.5 KCl, 26 NaHCO_3_, 1.5 MgCl_2_-6H_2_O, 1.3 NaH_2_PO_4_-H_2_O, 10 Glucose, 1.2 CaCl_2_ at 36 °C for 10 minutes. Slices were then recovered in the chamber at room temperature until use. The fluorescent dye TO-PRO-3 has previously been shown to robustly label pericytes in fixed tissue (Mai-Morente et al., 2021), and we have adapted this for imaging in acutely prepared live brain slices. Following recovery, slices were incubated in 1 µM TO-PRO-3 diluted in ACSF for 20 minutes at room temperature to label and identify the pericyte soma. All solutions used were continuously gassed with 95% O_2_ and 5% CO_2_.

### Pharmacology and ion substitution

All salts were obtained from Sigma-Aldrich. Please refer to the key resources table for sources and identifiers of reagents used. Time 0 represents the start of the first acquisition when the new solution is estimated to have reached the bath (estimated based on the flow rate). For Ca^2+^-free ACSF, CaCl_2_ was omitted from the ACSF and 2 mM EGTA was added. For 60 mM K^+^ ACSF, equimolar NaCl was replaced with KCl.

### Confocal imaging

Imaging was performed with a Zeiss LSM 510 laser scanning confocal microscope with a 40X water immersion objective lens (0.8NA). Pericytes in cortical brain slices were located by TO-PRO-3 and GCaMP6f co-localization and association with a Rhodamine B labelled vessel. GCaMP6f was excited with a 488 nm LED laser and was detected after passing through a 505-530 nm bandpass filter. TO-PRO-3 and Rhodamine B were excited with a 633 nm and 543 nm HeNe laser respectively, which were generally turned off during Ca^2+^ imaging. Ensheathing pericytes in a subset of explicitly labeled experiments were identified based on GCaMP fluorescence almost fully enwrapping the vessel, and were confirmed to be within 3 branch orders of the penetrating arteriole. Capillary pericytes of thin-stand or mesh morphology on ≥ 4 branch order were selected for all other experiments (no arteriole could be detected within 4 branches of the capillary, using the Rhodamine-IV signal). To measure calcium transient frequency over time, 50-second image acquisitions were made every 3-4 minutes in frame scanning mode at a 2 Hz sample rate. To record SOCE experiments and [Ca^2+^]_i_ rises evoked by ET-1 in Fig. 4, a single acquisition was made in frame scanning mode at a 2 Hz sample rate. Slices were perfused with ACSF at 2 mL/minute and were kept at 34 ± 2 °C during experiments.

### Data collection and analysis

Between frame XY-plane translational movement and within frame distortion were removed using a custom non-rigid alignment MATLAB algorithm, based on the *imregdemons* (AccumulatedFieldSmoothing = 2.5; PyramidLevels = 4; 32, 16, 8 and 4 iterations respectively for each pyramid level) and *imwarp* MATLAB functions (https://scanbox.org/2016/06/30/non-rigid-image-alignment-in-twenty-lines-of-matlab/). Hand-drawn regions of interest (ROI) separated pericyte somas from processes. The Astrocyte Quantification and Analysis (AQuA) MATLAB tool (Wang et al., 2019) was used for unbiased event-based Ca^2+^ signal analysis. A Gaussian filter was applied to the images (σ = 2), minimum event size was set to (pixels): 5/pixel size (µm), and detected events occurring in the same frame separated by (in pixels), 1/pixel size (µm) were merged. Event threshold parameters were set to reliably detect the majority of pericyte Ca^2+^ signaling events with minimal detection of false events outside the cell boundaries. Identical analysis parameters were used for all time series of a given cell and experiment. In experiments in which movies were collected at differing timepoints and average data was shown, a linear interpolation between data points was performed. Ca^2+^ transient data in Ca^2+^-free solution (Fig. 1) and after 1.2 mM Ca^2+^ washback (Fig. 1E) was analyzed from 5 minutes after solution was estimated to reach the bath (t=0) until cessation of treatment. Ca^2+^ transient data following agonist application (ET1 or U46619) in Ca^2+^ free solution (Fig, 1F,G) or GSK-7975A (Fig. 4F,G) was only transiently increased after agonist application before stores became depleted and was therefore analyzed over a brief period during which the agonists visibly evoked an increase in transient signals: ET-1 in Ca^2+^-free (65.2 ± 16.31 seconds); U46619 (78.86 ± 12.53 seconds); ET-1 in GSK-7975A (149 +/- 10.77 seconds). Ca^2+^ transient data during treatments in Figs. 2-3 was measured from 10-20 minutes after drug application unless otherwise stated. Transient frequency was normalized to 10 µm of pericyte processes (referred to as ‘length’). To measure SOCE and [Ca^2+^]_i_ rises evoked by ET-1 in Fig. 4, background fluorescence was subtracted, a hand-drawn ROI was placed around the soma, and Δ*F*/*F* was averaged 30 seconds around the peak value recorded. Event density heat maps in figures represent the fraction of frames in which an active event was present in each pixel.

### Statistics

For all statistical comparisons normality was first assessed with a Shapiro-Wilk normality test. Unless stated otherwise in figure legends, statistics were calculated with two-tailed paired Students’ t-tests. A Mann-Whitney U test was used to compare two groups when normality test failed.

### Software availability

All software used in the study is already available open source, for any additional information required please contact the corresponding author.

## Acknowledgements

This work was supported by an NSERC discovery grant (RGPIN-2020-05276) to RLR. RLR holds a Canada Research Chair in Neurovascular Interactions. BP was supported by an USRA award from NSERC and a CIHR MSc Canada Graduate Scholarship. The authors would like to thank Dr. Richard Robitaille (Université de Montréal) for gifting equipment, Dr. Terry Snutch (University of British Columbia) for gifting Z944, and Dr. Volkhard Linder (Maine Medical Center) for PDGFRβ-cre mice, and Dr. Isabel Laplante for colony and lab management.

## Competing interests

The authors declare no competing interests.

---

**Supplementary Figure 1:**
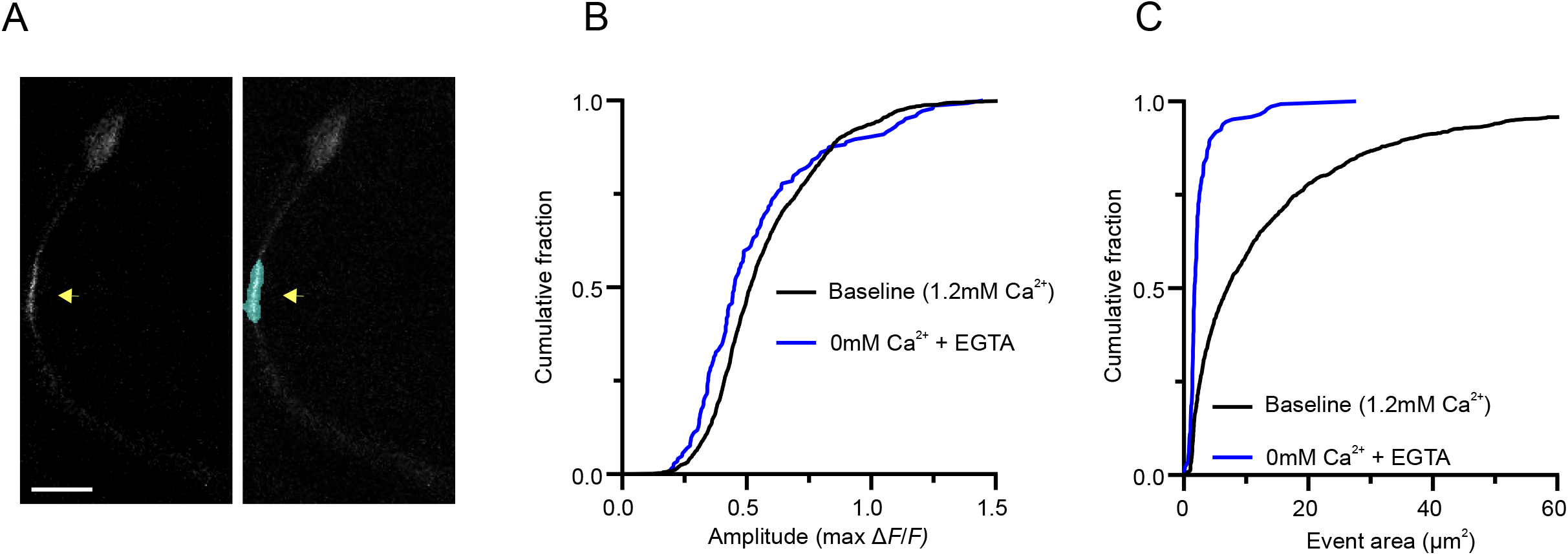
Related to Fig. 1, Imaging of pericyte microdomain Ca^2+^ signals and their dependence on extracellular Ca^2+^. **(A)** An example of an AQUA detected event in a capillary pericyte process. Scale bar = 10µm. **(B, C)** Cumulative histograms of event amplitude **(B)** and event area **(C)** at baseline and following wash out of extracellular Ca^2+^ (blue).

## Notes

### Competing Interest Statement

The authors have declared no competing interest.

## REFERENCES

Alarcon-Martinez L, Villafranca-Baughman D, Quintero H, Kacerovsky JB, Dotigny F, Murai KK, Prat A, Drapeau P, Polo AD. 2020. Interpericyte tunnelling nanotubes regulate neurovascular coupling. Nature 585:91–95. doi:10.1038/s41586-020-2589-x

Armulik A, Genové G, Betsholtz C. 2011. Pericytes: developmental, physiological, and pathological perspectives, problems, and promises. Developmental cell 21:193–215.

Armulik A, Genové G, Mäe M, Nisancioglu MH, Wallgard E, Niaudet C, He L, Norlin J, Lindblom P, Strittmatter K, Johansson BR, Betsholtz C. 2010. Pericytes regulate the blood-brain barrier. Nature 468:557–561. doi:10.1038/nature09522

Bell RD, Winkler EA, Sagare AP, Singh I, LaRue B, Deane R, Zlokovic BV. 2010. Pericytes Control Key Neurovascular Functions and Neuronal Phenotype in the Adult Brain and during Brain Aging. Neuron 68:409–427. doi:10.1016/j.neuron.2010.09.043

Chen BR, Kozberg MG, Bouchard MB, Shaik MA, Hillman EM. 2014. A critical role for the vascular endothelium in functional neurovascular coupling in the brain. Journal of the American Heart Association 3:e000787. doi:10.1161/JAHA.114.000787

Cuttler AS, LeClair RJ, Stohn JP, Wang Q, Sorenson CM, Liaw L, Lindner V. 2011. Characterization of Pdgfrb-Cre transgenic mice reveals reduction of ROSA26 reporter activity in remodeling arteries. Genesis 49:673–680. doi:10.1002/dvg.20769

Daneman R, Zhou L, Kebede AA, Barres BA. 2010. Pericytes are required for blood–brain barrier integrity during embryogenesis. Nature 468:562–566. doi:10.1038/nature09513

Derler I, Schindl R, Fritsch R, Heftberger P, Riedl MC, Begg M, House D, Romanin C. 2013. The action of selective CRAC channel blockers is affected by the Orai pore geometry. Cell Calcium 53:139–151. doi:10.1016/j.ceca.2012.11.005

Dias DO, Kalkitsas J, Kelahmetoglu Y, Estrada CP, Tatarishvili J, Holl D, Jansson L, Banitalebi S, Amiry-Moghaddam M, Ernst A, Huttner HB, Kokaia Z, Lindvall O, Brundin L, Frisén J, Göritz C. 2021. Pericyte-derived fibrotic scarring is conserved across diverse central nervous system lesions. Nat Commun 12:5501. doi:10.1038/s41467-021-25585-5

Gerhardt H, Betsholtz C. 2003. Endothelial-pericyte interactions in angiogenesis. Cell Tissue Res 314:15–23. doi:10.1007/s00441-003-0745-x

Glück C, Ferrari KD, Binini N, Keller A, Saab AS, Stobart JL, Weber B. 2021. Distinct signatures of calcium activity in brain mural cells. Elife 10:e70591. doi:10.7554/elife.70591

Gonzales AL, Klug NR, Moshkforoush A, Lee JC, Lee FK, Shui B, Tsoukias NM, Kotlikoff MI, Hill-Eubanks D, Nelson MT. 2020. Contractile pericytes determine the direction of blood flow at capillary junctions. Proc National Acad Sci 117:27022–27033. doi:10.1073/pnas.1922755117

Grant RI, Hartmann DA, Underly RG, Berthiaume A-AA, Bhat NR, Shih AY. 2017. Organizational hierarchy and structural diversity of microvascular pericytes in adult mouse cortex. Journal of cerebral blood flow and metabolism : official journal of the International Society of Cerebral Blood Flow and Metabolism 271678X17732229. doi:10.1177/0271678X17732229

Grubb S, Lauritzen M, Aalkjær C. 2021. Brain capillary pericytes and neurovascular coupling. Comp Biochem Physiology Part Mol Integr Physiology 254:110893. doi:10.1016/j.cbpa.2020.110893

Hariharan A, Robertson CD, Garcia DCG, Longden TA. 2022. Brain Capillary Pericytes are Metabolic Sentinels that Control Blood Flow through KATP Channel Activity. Biorxiv 2022.03.14.484304. doi:10.1101/2022.03.14.484304

Hariharan A, Weir N, Robertson C, He L, Betsholtz C, Longden TA. 2020. The Ion Channel and GPCR Toolkit of Brain Capillary Pericytes. Front Cell Neurosci 14:601324. doi:10.3389/fncel.2020.601324

Hartmann DA, Berthiaume A-A, Grant RI, Harrill SA, Koski T, Tieu T, McDowell KP, Faino AV, Kelly AL, Shih AY. 2021. Brain capillary pericytes exert a substantial but slow influence on blood flow. Nat Neurosci 24:633–645. doi:10.1038/s41593-020-00793-2

Hartmann DA, Coelho-Santos V, Shih AY. 2022. Pericyte Control of Blood Flow Across Microvascular Zones in the Central Nervous System. Annu Rev Physiol 84:331–354. doi:10.1146/annurev-physiol-061121-040127

Hill RA, Tong L, Yuan P, Murikinati S, Gupta S, Grutzendler J. 2015. Regional Blood Flow in the Normal and Ischemic Brain Is Controlled by Arteriolar Smooth Muscle Cell Contractility and Not by Capillary Pericytes. Neuron 87:95–110. doi:10.1016/j.neuron.2015.06.001

Hill-Eubanks DC, Werner ME, Heppner TJ, Nelson MT. 2011. Calcium Signaling in Smooth Muscle. Csh Perspect Biol 3:a004549. doi:10.1101/cshperspect.a004549

Iadecola C, Yang G, Ebner TJ, Chen G. 1997. Local and propagated vascular responses evoked by focal synaptic activity in cerebellar cortex. Journal of neurophysiology 78:651–9.

Jairaman A, McQuade A, Granzotto A, Kang YJ, Chadarevian JP, Gandhi S, Parker I, Smith I, Cho H, Sensi SL, Othy S, Blurton-Jones M, Cahalan MD. 2022. TREM2 regulates purinergic receptor-mediated calcium signaling and motility in human iPSC-derived microglia. Elife 11:e73021. doi:10.7554/elife.73021

Korte N, Ilkan Z, Pearson CL, Pfeiffer T, Singhal P, Rock JR, Sethi H, Gill D, Attwell D, Tammaro P. 2022. The Ca2+-gated channel TMEM16A amplifies capillary pericyte contraction and reduces cerebral blood flow after ischemia. J Clin Invest 132. doi:10.1172/jci154118

Lampl Y, Fleminger G, Gilad R, Galron R, Sarova-Pinhas I, Sokolovsky M. 1997. Endothelin in cerebrospinal fluid and plasma of patients in the early stage of ischemic stroke. Stroke J Cereb Circulation 28:1951–5. doi:10.1161/01.str.28.10.1951

Li Q, Puro DG. 2001. Adenosine activates ATP-sensitive K(+) currents in pericytes of rat retinal microvessels: role of A1 and A2a receptors. Brain Res 907:93–9. doi:10.1016/s0006-8993(01)02607-5

Longden TA, Dabertrand F, Koide M, Gonzales AL, Tykocki NR, Brayden JE, Hill-Eubanks D, Nelson MT. 2017. Capillary K+-sensing initiates retrograde hyperpolarization to increase local cerebral blood flow. Nature Neuroscience 20:717. doi:10.1038/nn.4533

Mai-Morente SP, Marset VM, Blanco F, Isasi EE, Abudara V. 2021. A nuclear fluorescent dye identifies pericytes at the neurovascular unit. J Neurochem 157:1377–1391. doi:10.1111/jnc.15193

Molnár T, Yarishkin O, Iuso A, Barabas P, Jones B, Marc RE, Phuong TTT, Križaj D. 2016. Store-Operated Calcium Entry in Müller Glia Is Controlled by Synergistic Activation of TRPC and Orai Channels. J Neurosci 36:3184–3198. doi:10.1523/jneurosci.4069-15.2016

Nakagomi T, Kubo S, Nakano-Doi A, Sakuma R, Lu S, Narita A, Kawahara M, Taguchi A, Matsuyama T. 2015. Brain Vascular Pericytes Following Ischemia Have Multipotential Stem Cell Activity to Differentiate Into Neural and Vascular Lineage Cells. Stem Cells 33:1962–1974. doi:10.1002/stem.1977

Nortley R, Korte N, Izquierdo P, Hirunpattarasilp C, Mishra A, Jaunmuktane Z, Kyrargyri V, Pfeiffer T, Khennouf L, Madry C, Gong H, Richard-Loendt A, Huang W, Saito T, Saido TC, Brandner S, Sethi H, Attwell D. 2019. Amyloid β oligomers constrict human capillaries in Alzheimer’s disease via signaling to pericytes. Science 365:eaav9518. doi:10.1126/science.aav9518

Ong HL, Souza LB de, Ambudkar IS. 2016. Role of TRPC Channels in Store-Operated Calcium Entry. Adv Exp Med Biol 898:87–109. doi:10.1007/978-3-319-26974-0_5

Prakriya M, Lewis RS. 2001. Potentiation and inhibition of Ca2+ release-activated Ca2+ channels by 2-aminoethyldiphenyl borate (2-APB) occurs independently of IP3 receptors. J Physiology 536:3–19. doi:10.1111/j.1469-7793.2001.t01-1-00003.x

Ratelade J, Klug NR, Lombardi D, Angelim MKSC, Dabertrand F, Domenga-Denier V, Salman RA-S, Smith C, Gerbeau J-F, Nelson MT, Joutel A. 2020. Reducing Hypermuscularization of the Transitional Segment Between Arterioles and Capillaries Protects Against Spontaneous Intracerebral Hemorrhage. Circulation 141:2078–2094. doi:10.1161/circulationaha.119.040963

Rungta RL, Chaigneau E, Osmanski B-F, Charpak S. 2018. Vascular Compartmentalization of Functional Hyperemia from the Synapse to the Pia. Neuron 99. doi:10.1016/j.neuron.2018.06.012

Rungta RL, Zuend M, Aydin A-K, Martineau É, Boido D, Weber B, Charpak S. 2021. Diversity of neurovascular coupling dynamics along vascular arbors in layer II/III somatosensory cortex. Commun Biology 4:855. doi:10.1038/s42003-021-02382-w

Rustenhoven J, Jansson D, Smyth LC, Dragunow M. 2017. Brain Pericytes As Mediators of Neuroinflammation. Trends Pharmacol Sci 38:291–304. doi:10.1016/j.tips.2016.12.001

Sancho M, Klug NR, Mughal A, Koide M, Cruz SH de la, Heppner TJ, Bonev AD, Hill-Eubanks D, Nelson MT. 2022. Adenosine signaling activates ATP-sensitive K+ channels in endothelial cells and pericytes in CNS capillaries. Sci Signal 15:eabl5405. doi:10.1126/scisignal.abl5405

Singh A, Hildebrand M, Garcia E, Snutch T. 2010. The transient receptor potential channel antagonist SKF96365 is a potent blocker of low-voltage-activated T-type calcium channels. Brit J Pharmacol 160:1464–1475. doi:10.1111/j.1476-5381.2010.00786.x

Vanlandewijck M, He L, Mäe M, Andrae J, Ando K, Gaudio F, Nahar K, Lebouvier T, Laviña B, Gouveia L, Sun Y, Raschperger E, Räsänen M, Zarb Y, Mochizuki N, Keller A, Lendahl U, Betsholtz C. 2018. A molecular atlas of cell types and zonation in the brain vasculature. Nature 554:475. doi:10.1038/nature25739

Várnai P, Hunyady L, Balla T. 2009. STIM and Orai: the long-awaited constituents of store-operated calcium entry. Trends Pharmacol Sci 30:118–128. doi:10.1016/j.tips.2008.11.005

Wang Y, DelRosso NV, Vaidyanathan TV, Cahill MK, Reitman ME, Pittolo S, Mi X, Yu G, Poskanzer KE. 2019. Accurate quantification of astrocyte and neurotransmitter fluorescence dynamics for single-cell and population-level physiology. Nat Neurosci 22:1936–1944. doi:10.1038/s41593-019-0492-2

Yang AC, Vest RT, Kern F, Lee DP, Agam M, Maat CA, Losada PM, Chen MB, Schaum N, Khoury N, Toland A, Calcuttawala K, Shin H, Pálovics R, Shin A, Wang EY, Luo J, Gate D, Schulz-Schaeffer WJ, Chu P, Siegenthaler JA, McNerney MW, Keller A, Wyss-Coray T. 2022. A human brain vascular atlas reveals diverse mediators of Alzheimer’s risk. Nature 603:885–892. doi:10.1038/s41586-021-04369-3

Yoast RE, Emrich SM, Zhang X, Xin P, Johnson MT, Fike AJ, Walter V, Hempel N, Yule DI, Sneyd J, Gill DL, Trebak M. 2020. The native ORAI channel trio underlies the diversity of Ca2+ signaling events. Nat Commun 11:2444. doi:10.1038/s41467-020-16232-6

